# Inconspicuous breeding coloration to conceal eggs during mouthbrooding in male cardinalfish

**DOI:** 10.1101/2024.05.07.592841

**Authors:** Hikaru Ishihara, Shinji Kanda

## Abstract

The body coloration of animals has various functions, such as camouflage coloration for hiding their existence from other organisms and conspicuous coloration for appealing to their potential mates. Since the breeding colorations of males are widely considered to attract females, most previous studies on breeding coloration have mostly focused on conspicuous breeding coloration, which may have prevented the discovery of possible functions other than appealing. Here, based on a speculation that such breeding coloration might occur in species with weak sexual selection in males, we focused on Apogoninae whose sex role is considered to be reversed due to their characteristic parental behavior, paternal mouthbrooding. Through detailed morphological observations, we found that males exhibit non-conspicuous white structures, consisting of iridophores, in the lower jaw during the breeding season. Artificial implantation of eggs inside the mouth in both sexes showed that the white structure in the lower jaws, which specifically exists in males, drastically reduces the visibility of eggs during mouthbrooding. This suggested that this coloration may serve to conceal the conspicuous coloration of eggs during mouthbrooding. In addition, *in vivo* and *in vitro* hormone treatment experiments revealed that iridophore development in the lower jaw is induced by androgen through the Alkal-Ltk pathway. These results suggest that androgen-dependent breeding colorations in males, which have been considered to attract females, may serve the opposite function in these species, “inconspicuous coloration”, to increase the fitness of their specialized behavior, mouthbrooding.

## Introduction

Animals exhibit various colorations, which play important roles in adapting to their environment. Most animals exhibit camouflage coloration to hide their existence from predators or prey. Among them, countershading, which is characterized by darker coloring on the dorsum than on the ventrum, is one of the most conserved colorations and is considered an adaptation for camouflage in the natural environment^1,2^. On the other hand, many animals sometimes exhibit conspicuous coloration as a signal for other individuals. Breeding season-specific coloration, called breeding coloration or nuptial coloration, is recognized to serve as conspicuous signals for courtship with the opposite sex or agonistic interactions among individuals of the same sex^3^. This trait usually evolved specifically in males who have been strongly sexually selected.

Teleosts have been served as excellent models for studying the mechanisms and significance of male-specific breeding colorations. For instance, male three-spined stickleback (*Gasterosteus aculeatus*)^4^ and male guppies (*Poecilia reticulata*)^5,6^ are well known to exhibit conspicuous coloration during the breeding season for mating success. Importantly, these male-specific breeding colorations are mostly induced by androgens^7–9^, sex steroids that are important for male functions. On microscopic levels, several studies have revealed that androgens induce chromatophore development such as melanophores, xanthophores, erythrophores, iridophores, and leucophores^10–13^ to form male-specific breeding colorations. On the other hand, the molecular mechanisms underlying this phenomenon have been partially analyzed in melanophores^12^ and xanthophores^12,14,15^, whereas those of others are unclear to date.

Although the functions of body coloration are not limited to appealing to other individuals, previous studies of male-specific coloration have concentrated on phenomena involving conspicuous breeding coloration. Therefore, identification of non-conspicuous color changes in males during the breeding season, if present, might be a missing piece toward understanding the functions and mechanisms of male-specific breeding coloration. Here, we speculated that such non-conspicuous male-specific coloration might be found in species with weak sexual selection in males. Specifically, we focused on the body coloration of Apogoninae, in which the sex role is often reversed, and the chooser sex is male due to their characteristic parental behavior, paternal mouthbrooding, which imposes a high cost to males^16–19^. In several species from two genera in this subfamily, we found that male fish have a white lower jaw where females show a transparent structure, although this difference is ambiguous when the mouth is closed (spotnape cardinalfish (*Ostorhinchus notatus*), *Ostorhinchus doederleini*, *Ostorhinchus semilineatus*, and *Sphaeramia nematoptera*). Given the characteristic parental behavior in these genera, we hypothesized that this non-conspicuous coloration in the mouthes might have a function during mouthbrooding.

In the present study, to understand the possible functions and mechanism of non-conspicuous breeding coloration, we investigated a male-specific coloration of the lower jaw in an Apogoninae, spotnape cardinalfish as a model. First, we found that the lower jaws of males but not females have a white structure, which drastically reduces the visibility of the conspicuous coloration of eggs inside the mouth. We also found that this male-specific white structure consists of iridophores, which can conceal the eggs inside their mouth and maintain the countershading pattern of the parental fish. Furthermore, we revealed that this coloration is season- and male-specific and is induced by androgens, which implies that common mechanism for conspicuous breeding colorations are repurposed to conceal eggs during mouthbrooding in these species. Additionally, we demonstrated that this androgen-induced iridophore development is mediated by the activation of the Alkal-Ltk pathway. These results provide functional implication of non-conspicuous breeding coloration, which may extend the possible biological significance of male-specific breeding colorations.

## Results

### Male spotnape cardinalfish have whiter lower jaws

To investigate sexually dimorphic coloration in spotnape cardinalfish, we observed the gross morphology of sexually mature spotnape cardinalfish. There was no apparent difference between females and males (Figure 1A). However, upon opening the mouth, we observed sexual dimorphism in a region of the lower jaw that is likely to be more exposed during mouthbrooding in males. Here, only males exhibited white connective tissues and musculus intermandibularis in the lower jaw, while females possessed more transparent structures (Figure 1B Open). Note that this sexually dimorphic coloration was ambiguous when the mouth was closed (Figure 1B Close). Furthermore, we found that this dimorphism is conserved in other mouthbrooding cardinalfishes from two genera in Apogoninae: *Sphaeramia nematoptera*, *Ostorhinchus doederleini*, and *Ostorhinchus semilineatus* (Figures S2A-C). This observation was consistent with a recently reported sexually dimorphic morphology of *Ostorhinchus cyanosoma*, which was used to distinguish sex in the aquaculture field^20^.

**Figure 1.**
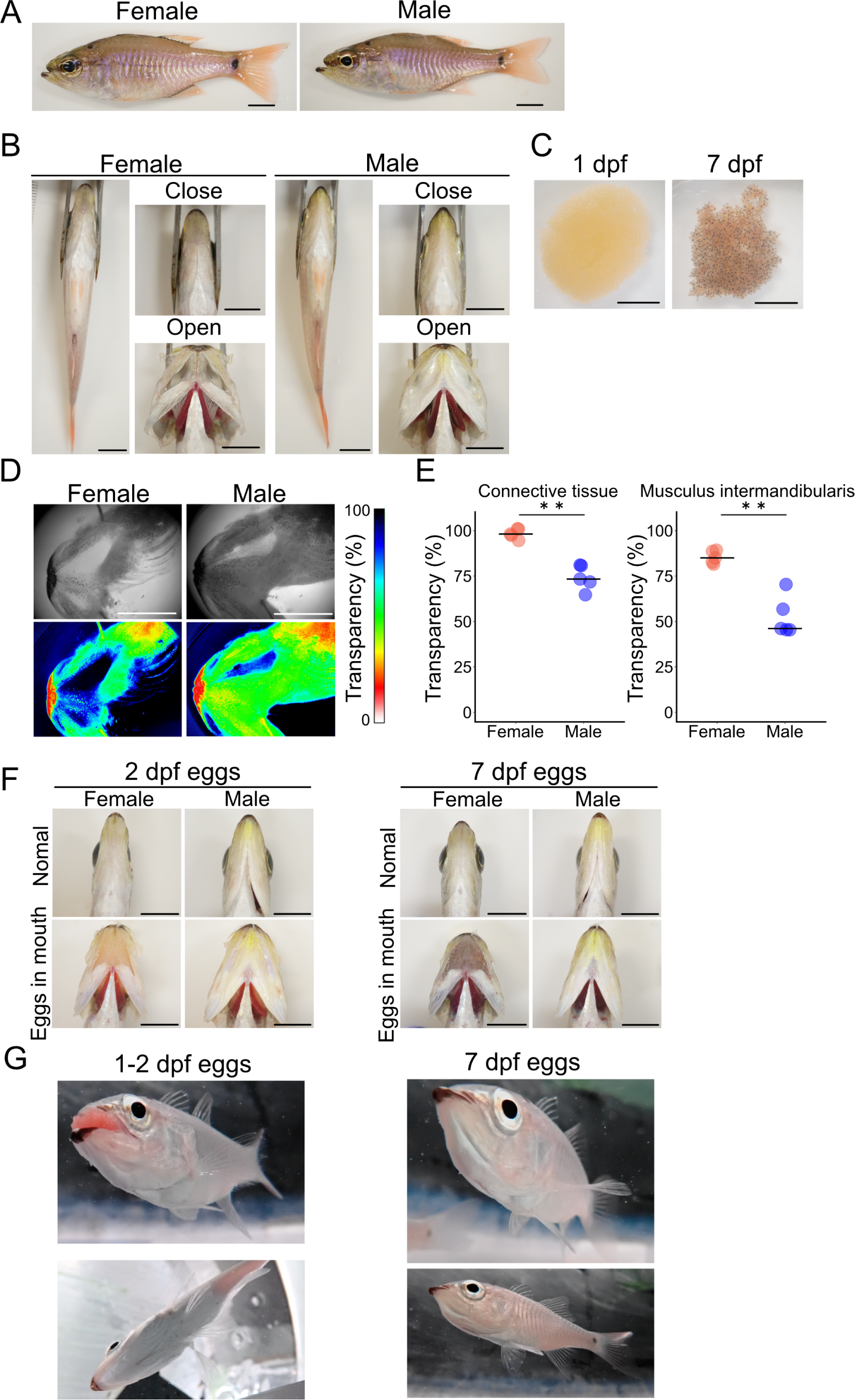
Male spotnape cardinalfish have a whiter lower jaw during mouthbrooding, which conceals their eggs. (A) Whole-body photograph of a mature spotnape cardinalfish. There is no apparent difference between female and male. (B) The ventral region and lower jaw of mature spotnape cardinalfish. When the mouth is closed, there is no major difference between the male and female lower jaw. However, when the mouth is opened, males exhibit a white lower jaw while females show transparent one. (C) Egg masses taken from the male mouth. At 1 days post fertilization (dpf), the coloration of the eggs is orange, whereas the coloration ranges from orange to black at 7 dpf. (D) Representative images showing transparency in the lower jaw of a mature female and male. Transparency is show in a pseudo-color. (E) The transparency of the connective tissue and musculus intermandibularis in the lower jaw is significantly lower in males than in females. The bars indicate the medians. **, p < 0.01, Wilcoxon rank sum test, n = 5. (F) Photographs showing mouths artificially filled with developing eggs. Only the male mouths are not transparent and maintain the whiteness of their ventral region. (G) Underwater photographs of mature males brooding eggs showing that egg coloration are not reflected to the surface of the lower jaw and that the ventral region remains whitish during mouthbrooding. Scale bars, 10 mm. See also Figures S1A-C showing the lower jaw area measured in the transparency analysis, and Figure S2.

### The whiteness and opaqueness of the male lower jaw may serve to conceal the coloration of eggs

Due to the their reversed sex roles, we consider that this coloration in males should not serve as a conspicuous signal for intersexual or intrasexual competition. Instead, we hypothesized that this male-specific coloration may conceal the conspicuous colorations of eggs inside the mouth during mouthbrooding. First, we observed the coloration of eggs at different developmental stages. The coloration of the eggs was orange at 1 day post fertilization (dpf), while ranged from orange to black at 7 dpf (Figure 1C). Given that the ventral surface, including the lower jaw, is generally whitish to maintain the countershading pattern, these egg colorations may disrupt the countershading pattern during mouthbrooding if the lower jaw is transparent. Next, to test whether the male-specific whitish lower jaw can serve to conceal the coloration of eggs, we compared the transparency of the lower jaws of sexually mature males and females. The transparencies of the connective tissue and musculus intermandibularis in the lower jaw were significantly lower in males than in females (Figures 1D and 1E). Finally, to investigate whether the whitish lower jaw of males serves to conceal the coloration of eggs, we artificially introduced the developing eggs into sexually mature female and male mouths. When we introduced eggs into the female mouth, the orange to black coloration of the eggs was obviously reflected to the surface (Figure 1F). On the other hand, the coloration of eggs was not reflected to the surface, and the ventral region remained whitish when we introduced eggs into the male mouth. These results indicate that the whitish lower jaw of males serves to conceal the coloration of eggs during mouthbrooding. Additionally, we observed male fish spontaneously started mouthbrooding to confirm that the coloration of the eggs are concealed (Figure 1G). The lower jaw of the mouthbrooding male was swollen due to the presence of eggs inside its mouth. On the other hand, the coloration of the lower jaw remained whitish despite the egg coloration, which has been suggested from the above observations.

These results suggest that the whiteness and opaqueness of the male lower jaw serve to conceal the coloration of eggs, which may maintain countershading during mouthbrooding. This camouflage coloration may reduce the possibility of male parental fish being found by other organisms, such as predators.

### Iridophores contribute to male-specific whitish coloration in the lower jaw

Sexually dimorphic coloration often consists of chromatophores such as xanthophores, melanophores, and iridophores. Given this background, we tried to identify what type of cells contribute to male-specific whitish and opaque coloration in the lower jaw. Observation under a stereoscopic microscope revealed that there are many irregularly shaped cells in the male connective tissue in the lower jaw. These cells showed a property of being highly visible under reflected light but ambiguous under transmitted light (Figure 2A), which is characteristic of iridophores. For more detailed observation, we conducted a histological analysis of the lower jaw using hematoxylin and eosin (HE)-stained sections. Specifically, in males, a slightly dark region was observed in the epithelium of the lower jaw (Figure 2B Bright field). Since iridophores have birefringent properties under polarized illumination^21,22^, we observed tissue sections under polarized illumination. The dark region observed in the bright field was bright white‒green under polarized illumination (Figure 2B, The observed region is illustrated in Figure S3). In addition, by *in situ* hybridization, the expression of the *purine nucleoside phosphorylase 4a* (*pnp4a*), known as an iridophore marker in zebrafish (*Danio rerio*)^23^, and Japanese medaka (*Oryzias latipes*)^24^, which is involved in the synthesis of guanine for reflecting platelets in iridophores, was detected in the same region (Figure 2C). Furthermore, qRT‒ PCR analysis revealed that the expression level of the iridophore marker gene *pnp4a* was significantly greater in males than in females in connective tissues and in musculus intermandibularis in the lower jaw, whereas it was not significantly different in other tissues (Figure 2D). These results revealed that male-specific iridophores in the epithelium contribute to male-specific whitish coloration in the lower jaw.

**Figure 2.**
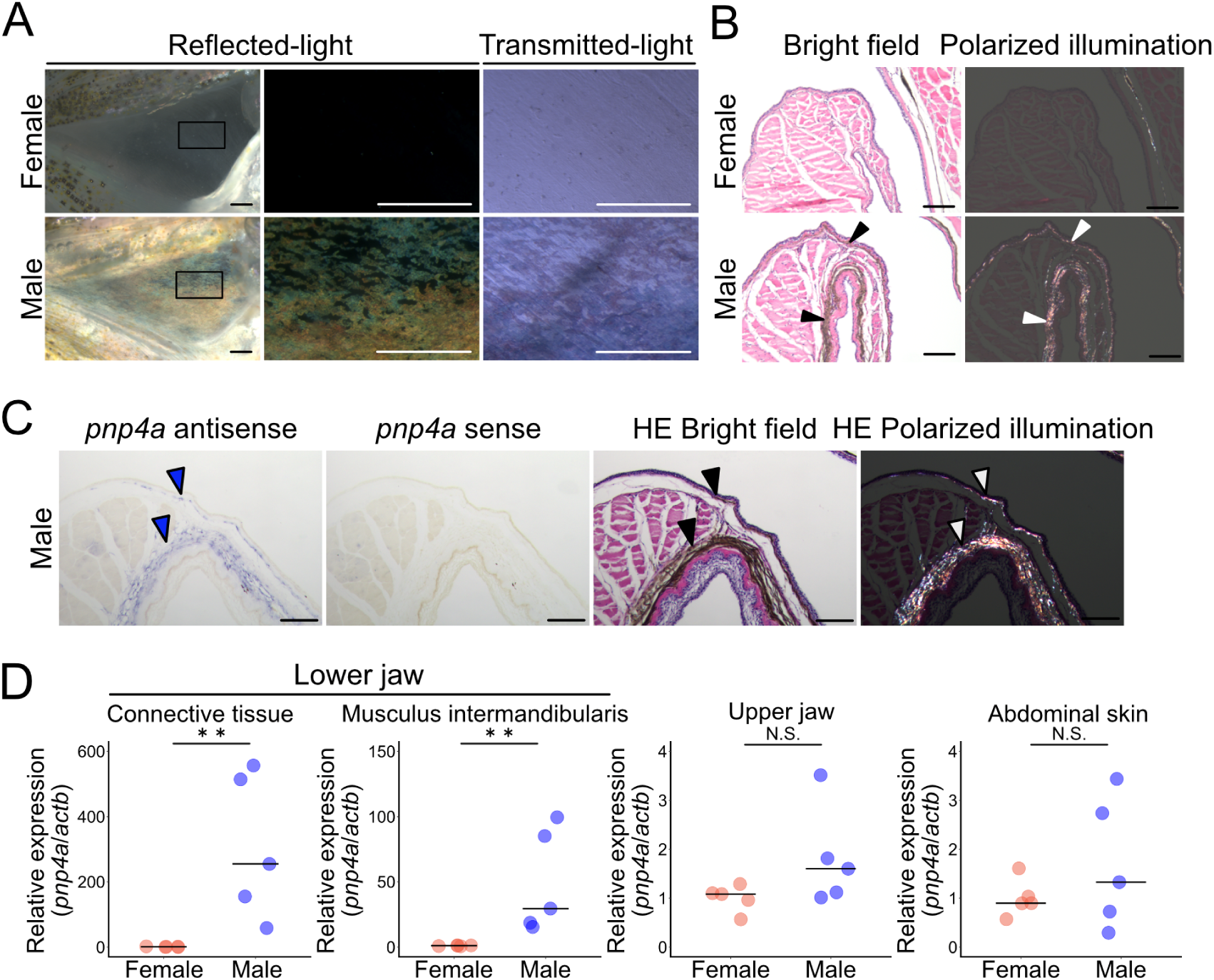
Male-specific development of iridophores in the lower jaw causes whiteness in the breeding season. (A) The appearance of the white structure of the lower jaw in males under a stereoscopic microscope resembles the characteristics of iridophores. Under reflected light, many irregularly shaped cells become highly visible, whereas under transmitted light, the cells in the lower jaw of males are ambiguous. The square frames in the left photographs indicate the locations of the enlarged views shown in the right photographs. Scale bar, 2 mm. (B) Hematoxylin and eosin (HE)-stained frontal section of the lower jaw of female and male. Under the bright field, dark regions are observed in the epithelium of the lower jaw, specifically in males (black arrowheads). Under polarized illumination, the dark region in the bright field is bright white‒green due to the birefringent properties (white arrowheads). Scale bar, 100 μm. (C) *In situ* hybridization of an iridophore marker gene, *purine nucleoside phosphorylase 4a* (*pnp4a*) and hematoxylin and eosin (HE)-stained sections of the lower jaw of male. The signals of *pnp4a* (blue arrowhead) were observed to correspond to the dark region in HE-stained sections under brightfield microscopy (black arrowhead) and the bright white‒green region under polarized illumination (white arrowhead). These observations indicates that the lower jaw of males but not females possess iridophores. Scale bar, 100 μm. Note that no purple signals are detected with the control sense probe. (D) The expression level of *pnp4a*, of males is significantly greater than that of females in both connective tissue and musculus intermandibularis in the lower jaw. On the other hand, there were no significant differences in other tissues, such as the upper jaw and abdominal skin. The bars indicate the medians. **, p < 0.01, N.S., not significant, Wilcoxon rank sum test; n = 5. See also Figures S1A-C showing the descriptions of the lower jaw area used in the observation, and Figure S3.

### Androgens induce male-specific whitish coloration in the lower jaw as a breeding coloration

Since male-specific colorations often develop in sexually mature fish during the breeding season, we investigated the developmental timing and seasonal changes in the formation of male-specific whitish coloration in the lower jaw. First, we found that unlike adults, sexually immature males as well as females do not exhibit a whitish lower jaw (Figure 3A). Next, we investigated possible seasonal changes in the coloration of the lower jaw. Since the breeding season for this species is reported to be summer^25^ (June to October), we obtained wild spotnape cardinalfish in March, April, and July to observe the lower jaw. Notably, the gonad somatic index (GSI) increased from March to June (Figure S4), as previously reported. Here, we found that iridophores in the male lower jaw increase as the breeding season approaches, while they do not change in females (Figures 3B and 3C, Note that statistical analysis was not performed in March due to the small number of samples.). These results indicate that male-specific whitish coloration in the lower jaw develops along with sexual maturation in the breeding season.

**Figure 3.**
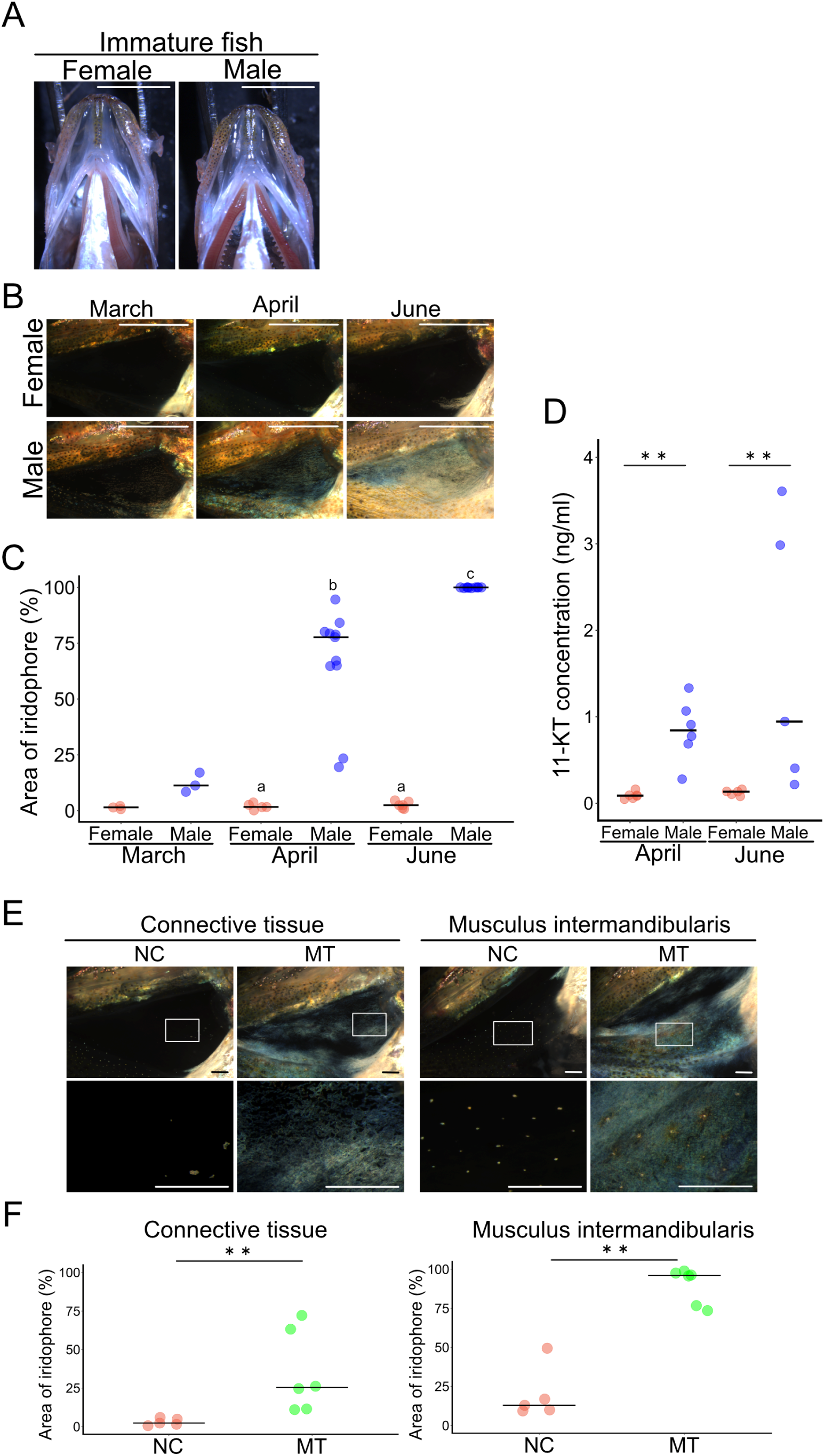
Androgens induce iridophore development in the lower jaw. (A) Immature female and male fish do not exhibit iridophores in the lower jaw. Scale bar, 5 mm. (B) Representative images showing that the number of iridophores in the connective tissue in the lower jaws of males increases as the breeding season approaches (from March to June). On the other hand, iridophores in the connective tissue in the lower jaw of females do not change as the breeding season approaches (from March to June). (C) The area of iridophores in the connective tissue in the lower jaw of males increases as the breeding season approaches (from March to June). The bars indicate the medians. Different letters indicate statistically significant differences (p < 0.05, Steel–Dwass test); females and males in March: n = 3; females in April: n = 5; males in April: n = 11; females in June: n = 6; males in June: n = 10. Note that statistical analysis was not performed in March due to the small number of samples. (D) The serum 11-ketotestosterone (11-KT) level is significantly greater in males than in females from prebreeding (April) to the breeding season (June). Bars indicate the median. **, p < 0.01, Wilcoxon rank sum test; females and males in April: n = 6; females and males in June: n = 5. (E) Representative images showing that there are no iridophores in the connective tissue or musculus intermandibularis in the lower jaw of EtOH-treated females (NC), whereas there are iridophores in the connective tissue and musculus intermandibularis in the lower jaw of 33 nM methyl testosterone-treated females (MT). The square frames in the upper photographs indicate the locations of the enlarged views shown in the lower photographs. Scale bar, 2 mm. (F) The area of iridophores in the connective tissue and musculus intermandibularis in the lower jaw of 33 nM methyl testosterone-treated females (MT) is significantly larger than that of EtOH-treated females (NC). The bars indicate the medians. **, p < 0.01, Wilcoxon rank sum test; NC: n = 5; MT: n = 6. See also Figure S1 showing the lower jaw area used to measure the area of the iridophore and for the observations, Figures S4 and S5.

The coloration specifically developed during the breeding season, known as breeding coloration, is often dependent on androgens. Therefore, we hypothesized that androgens may induce iridophore development in the lower jaw. In fact, consistent with the development of coloration, the plasma concentration of a potent androgen in teleosts, 11-ketotestosterone (11-KT), was significantly greater in males than in females in April and June (Figure 3D). In addition, the expression of *androgen receptor a* (*ara*) was detected, and *androgen receptor b* (*arb*) was weakly detected in the connective tissue and musculus intermandibularis of the male lower jaw (Figure S5A). These results suggested that androgens may induce iridophore development in the lower jaw.

To test this possibility, we exposed spotnape cardinalfish to methyl testosterone (MT) and observed their lower jaws. Here, we used females because they do not spontaneously accumulate iridophores in their lower jaws. Notably, females are assumed to be receptive to androgens, as the expression levels of *ara* and *arb* in the lower jaw were comparable to those in males (Figure S5B). As we expected, female fish treated with MT for 16 days exhibited a significantly larger iridophore area than female fish treated with EtOH as a solvent control in the connective tissue and musculus intermandibularis in the lower jaw (Figures 3E and 3F). These findings strongly suggested that androgens induce iridophore development in the lower jaw.

Taken together, these results indicated that androgens induce iridophore development, which induces whiteness in the lower jaw in sexually mature males during the breeding season.

### Androgens activates the iridophore development pathway

To explore the molecular mechanisms underlying androgen-dependent iridophore development in the lower jaw, we performed a transcriptome analysis on the musculus intermandibularis in the lower jaw of female fish treated with MT as a model. First, we identified the differentially expressed genes (DEGs) between female fish treated with MT or EtOH for 7 days, corresponding to the time when iridophores appeared in the lower jaw in MT-treated females. We identified 165 upregulated DEGs by 7 days of MT treatment (Figure 4A). These genes include several known iridophore-related genes, such as the iridophore marker gene *pnp4a*, *alk and ltk ligand 2a* (*alkal2a*)^26,27^, *four-and-half LIM domain transcription factor 2b* (*fhl2b*)^28,29^, *apolipoprotein Da1* (*apoDa1*)^28^, *glycoprotein nmb* (*gpnmb*)^28^, and *endothelin receptor ba* (*ednrba*)^30,31^. Next, to investigate whether these iridophore-related genes are upregulated by androgen prior to iridophore development and can serve as triggers of iridophore development, we performed a similar experiment examining the effects of MT treatment for 3 days, which is insufficient for the emergence of iridophores, while prior processes are expected to progress. While 10 genes were detected as upregulated DEGs, none of them were likely related to iridophore functions (Figure S6). Therefore, we analyzed the data from the 3 day MT treatment experiment, focusing on iridophore-related genes that had been detected as upregulated DEGs after 7 days of MT treatment. None of these genes showed significant changes in expression levels. However, we found that the average expression of *alkal2a*, which was not detected as a DEG due to its low expression and small sample size, was approximately three times greater in the fish treated with MT than in those treated with EtOH (Figure 4B).

**Figure 4.**
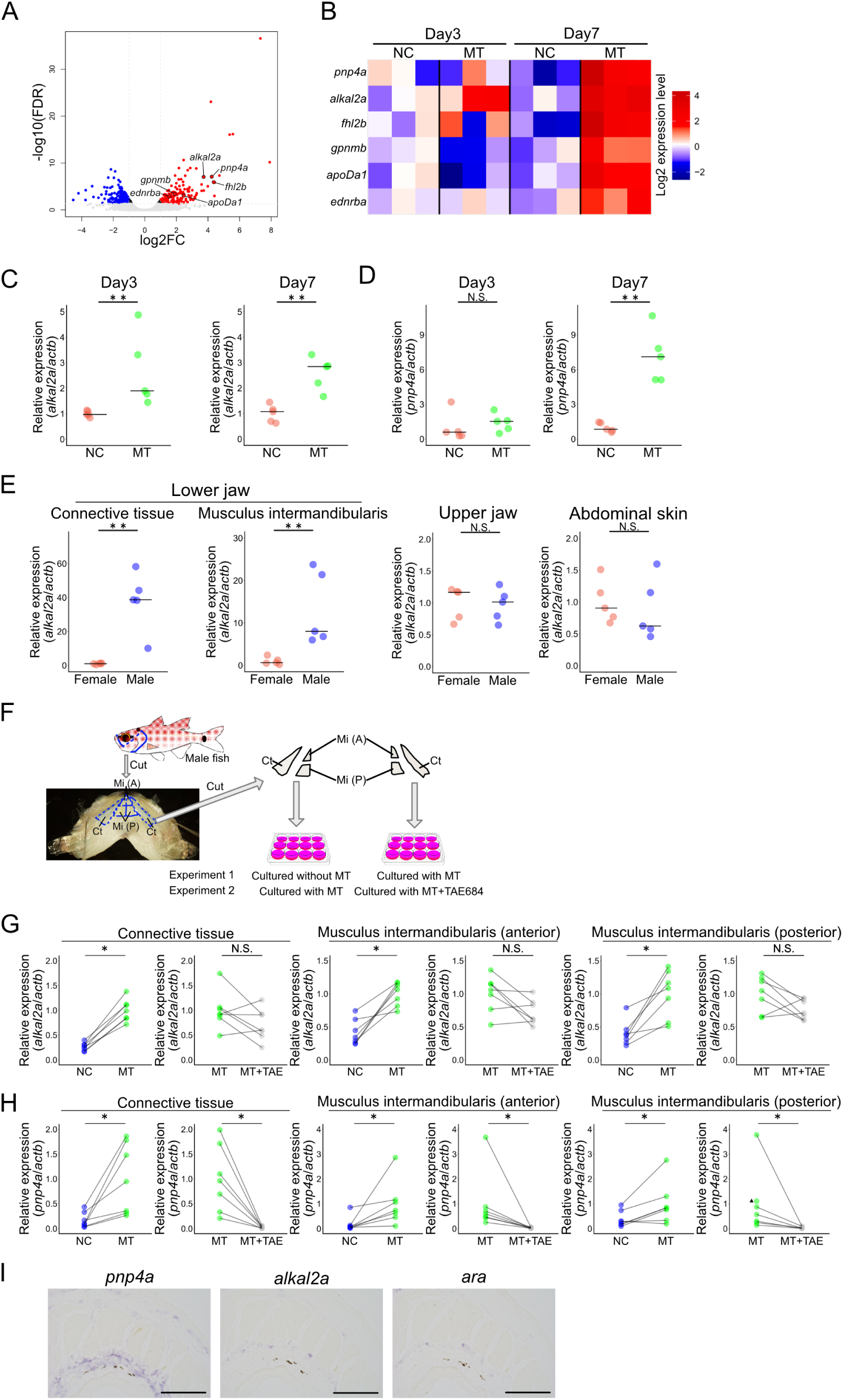
The iridophore development pathway is activated by androgen treatment. (A) Volcano plot of differentially expressed genes (DEGs) from RNA-seq analysis of musculus intermandibularis in the lower jaw between EtOH-treated females and 33 nM methyl testosterone-treated females on day 7. Positive log2 fold-change (FC) values correspond to increased expression in females treated with 33 nM methyl testosterone. The vertical lines indicate a log2FC threshold of 1, while the horizontal line indicates a -log10 false discovery rate (FDR) threshold of -log10(0.05). (B) Heatmap of the gene expression levels detected by RNA-seq analysis on treatment days 3 and 7 for 6 iridophore-related genes detected in the day 7 DEG analysis. The color represents the log2 expression level calculated with the mean expression level of EtOH-treated females (NC) on day 3 as 1. (C) The expression level of *alk and ltk ligand 2a* (*alkal2a*) of 33 nM methyl testosterone-treated females (MT) on day 3 and day 7 is significantly greater than those of EtOH-treated females (NC) in musculus intermandibularis in the lower jaw. Bars indicate the medians. **, p < 0.01, Wilcoxon rank sum test; n = 5. (D) The expression level of *pnp4a* of 33 nM methyl testosterone-treated females (MT) on day 7 is significantly greater than that of EtOH-treated females (NC) in musculus intermandibularis in the lower jaw, whereas it was not significantly different on day 3. The bars indicate the medians. **, p < 0.01, N.S., not significant, Wilcoxon rank sum test; n = 5. (E) The expression level of *alkal2a* in both the connective tissue and musculus intermandibularis in the lower jaw is significantly greater in males than in females, whereas it was not different in other tissues, such as the upper jaw and abdominal skin. The bars indicate the medians. **, p < 0.01, N.S., not significant, Wilcoxon rank sum test; n = 5. (F) Schematic illustration of *in vitro* culture of the lower jaw. The lower jaws were cut by the blue dashed line on both sides of the body in the fish illustration. Then, the connective tissues and anterior and posterior regions of the musculus intermandibularis were cut out by the blue dashed line. Note that each tissue sample was cut along the midline to obtain symmetrical tissues on the left and right sides. (G) The expression level of *pnp4a* in connective tissue and musculus intermandibularis in the lower jaw subjected to in vitro culture. *pnp4a* expression is greater in the sample cultured in 33 nM methyl testosterone (MT) than in the sample cultured in DMSO (NC). In the presence of 33 nM MT, the expression of *pnp4a* in the connective tissue of the lower jaw is significantly suppressed by the Alkal2a receptor Ltk inhibitor TAE684 (100 nM). The gray lines connect samples that originated from the same fish. *, p < 0.05, N.S., not significant, Wilcoxon signed-rank test; n = 7. (H) The expression level of *alkal2a* in connective tissue and musculus intermandibularis in the lower jaw subjected to in vitro culture. *alkal2a* expression was greater in the sample cultured in 33 nM methyl testosterone (MT) than in the sample cultured in DMSO (NC). In the presence of 33 nM MT, there is no significant difference between tissued cultured with Alkal2a receptor Ltk inhibitor TAE684 (100 nM) and without it. The gray lines connect samples that originated from the same fish. *, p < 0.05, N.S., not significant, Wilcoxon signed-rank test; n = 7. The black triangle represents a sample of fish tissue treated with MT, whose corresponding tissue exposed to MT+TAE684 showed an expression level below the detection limit. Therefore, no line or dot in MT+TAE was drawn. (I) *In situ* hybridization of adjacent sections suggested that a subset of iridophores expresses *alkal2a* and *ara*. The expressions of *alkal2a* and *ara* are detected in the inner regions of the area expressing the iridophore marker gene *pnp4a*. Note that the dark dot in each section consists of other types of chromatophores rather than signals. Scale bar, 100 μm. See also Figures S6, S7, and S8.

To further examine these results and the observed tendencies, we conducted another study of MT treatment to female fish and assessed the expression levels of *alkal2a* and *pnp4a* by RT‒qPCR over the same timescale as transcriptome analysis with a larger number of samples. In the musculus intermandibularis, the expression level of *alkal2a* was significantly greater in fish treated with MT for 3 and 7 days than in fish treated with EtOH (Figure 4C). On the other hand, the expression level of *pnp4a*, which is necessary for iridophore function, did not significantly differ between fish treated with MT and those treated with EtOH for 3 days. However, it was significantly geater in the fish treated with MT for 7 days than in those treated with EtOH (Figure 4D), which was consistent with the transcriptome analysis. Moreover, this trend was also observed in the connective tissue in the lower jaw, whereas it was not observed in the abdominal skin, which constitutively contains iridophores (Figure S7). In agreement with these results, in intact fish, the expression level of *alkal2a* was significantly greater in males than in females in the connective tissues and musculus intermandibularis in the lower jaw, whereas it was not significantly different in other tissues (Figure 4E). It was reported that *alkal2a* encodes a protein that is known to induce iridophore proliferation and differentiation during normal body color formation in embryonic and adult zebrafish in an autocrine and/or paracrine manner to activate its receptor, leukocyte tyrosine kinase (Ltk)^26,27^. It should be noted that the high sequence homology of the FAM150 domain and the conservation of four cysteines at the C-terminus in vertebrates, including spotnape cardinalfish, support the functional conservation of this identified *alkal2a* gene in terms of its activation of the receptor Ltk (Figure S8). These results suggested that *alkal2a* is upregulated by androgens and may trigger iridophore development.

To test this possibility, we treated tissues with an inhibitor during androgen-induced iridophore development in an isolated culture of the lower jaw (the experimental scheme is shown in Figure 4F). TAE684 is known to be a specific inhibitor of Ltk and its family gene Anaplastic lymphoma kinase (Alk) in mammals, and its activity has also been confirmed in zebrafish^32–34^. First, we compared the expression levels of *alkal2a* as well as *pnp4a*, an iridophore marker gene, between tissues cultured with DMSO (as a solvent control) and those cultured with MT. The expression levels of *alkal2a* and *pnp4a* were significantly greater in tissues cultured with MT than in those cultured with DMSO (Figures 4G and 4H). Next, we compared the expression levels of these genes between tissues cultured with MT alone and tissues cultured with MT in combination with the Ltk inhibitor TAE684. While the expression level of *alkal2a* was not significantly different between these two conditions, the expression level of *pnp4a* was significantly lower in tissues cultured with MT and TAE684 than in those cultured with MT. Here, since the inhibition of Ltk during androgen-treatment drastically reduced the expression of the iridophore marker *pnp4a*, androgen-induced iridophore development was suggested to be mediated by Ltk. Moreover, given that the expression of the Ltk ligand Alkal2a is increased by androgen treatment, Alkal2a may be the key molecule of this pathway.

Additionally, to investigate which type of cells receive androgens and express *alkal2a*, we conducted *in situ* hybridization for *alkal2a*, *ara* (the gene encoding the main androgen receptor in the lower jaw of spotnape cardinalfish, as shown in Figure S5A), and the iridophore marker gene *pnp4a* using adjacent lower jaw sections. The signals of *ara* and *alkal2a* were detected in the same region, which is located on the inner side of the area expressing *pnp4a* (Figure 4I, The observed region is illustrated in Figure S3). This finding is consistent with a previous report suggesting that iridophores express *alkal* in zebrafish embryos^26^. This result suggested that a subset of iridophores located on the inner side receive androgens and secrete Alkal2a in an autocrine and/or paracrine manner, which activates Ltk and promotes iridophore development.

## Discussion

This study showed that male spotnape cardinalfish have a whiter and opaque lower jaw, which may serve to conceal the conspicuous coloration of eggs. This male-specific coloration was proven to be caused by iridophores that are induced by androgens during the breeding season. Additionally, we indicated that androgen-dependent iridophore development is mediated by the Alkal2a-Ltk pathway. Although breeding colorations induced by androgens are often known as conspicuous coloration, our data suggest that cardinalfishes repurpose them as camouflage coloration during mouthbrooding, thereby extending the biological significance of male-specific breeding colorations.

We suggested that the whiteness and opaqueness of the male lower jaw may conceal the conspicuous coloration of the eggs (Figures 1D-G). To date, almost all male-specific breeding coloration has been considered to serve as a conspicuous signal to other individuals of the same species and has evolved through sexual selection. For instance, the male threespine stickleback has a red throat coloration, which serves as both a cue to attract a potential female mate and a threat signal in male‒ male competition^4,35,36^. However, it is believed that the sex role is reversed in Apogoninae, and choosier sex is considered to be male due to its costly parental behavior^16–19^. Therefore, it is unlikely that the male-specific coloration commonly observed in this subfamily serves to attract females. It is important to note that we observed a whiter lower jaw in males of another Apogoninae, *Ostorhinchus doederleini*, whose sexual role has been reported to be reversed^16^ (Figure S2B), although the occurrence of this sex role reversal is still controversial in spotnape cardinalfish^37–41^. Besides, we observed that the mating behavior of spotnape cardinalfish is obviously more active and aggressive in females than in males (Figure S10). Furthermore, this coloration is unlikely to serve as a conspicuous signal because most of this sexually dimorphic region is usually folded and is rarely exposed except for during mouthbrooding (Figure 1B). For these reasons, we considered that the whitish coloration of the male lower jaw serves as camouflage coloration during mouthbrooding, rather than as a signal to other individuals. Concealing colorful eggs may be advantageous as it helps maintain countershading during mouthbrooding as clearly shown in the sexually different appearance when eggs are implanted in the mouth (Figure 1F). Countershading, characterized by darker coloring on the dorsum and lighter coloring on the ventrum, is a widespread form of camouflage that reduces the risk of detection by predators. Given the coloration of the eggs from orange to black (Fig. 1C), male spotnape cardinalfish require the structure to conceal the egg coloration to maintain a whitish ventrum during mouthbrooding to reduce their predation risk. If not, the coloration of the orange to black eggs is directly reflected to the ventral surface of the parental fish to disrupt the countershading pattern (Figure 1F). Although there is room to consider whether this coloration might play a role in intraspecies relationships, the seasonal and sex specificity of this coloration as well as the gross morphology of jaws and eggs strongly suggest that this coloration plays an important role in concealing eggs. Also, given this function, it can be concluded that this phenomenon has been acquired not by sexual selection pressure, but by predation pressure.

We also identified that this breeding male-specific coloration is due to the iridophore development in breeding conditions (Figures 2, 3B, and 3C). As described above, this coloration can be regarded as an inconspicuous breeding coloration, unlike many well-known conspicuous breeding colorations reported so far (e.g. male three-spined stickleback^4^; male guppies^5,6^). Interestingly, contrary to this difference in the functions or appearance, the mechanism underlying the development of this inconspicuous structure was suggested to be similar to what develops conspicuous breeding colorations, especially in terms of androgen dependence (Figures 3). Thus, they may have repurposed this androgen-dependent iridophore development for hiding their eggs after the acquisition of the mouthbrooding trait. Interestingly, although different from mouthbrooding, a similar phenomenon was identified in eels^42^. Eels show silvering during breeding migration to the sea, which includes a body-color change to form countershading in the open sea^42^. Despite being observed in both males and females, this change is also suggested to be androgen-dependent^43,44^. Therefore, there may be a common mechanism by which androgens can change body coloration during the breeding season, while selection pressure can lead to either conspicuous or camouflage coloration according to the situation and tissue.

While many fish exhibit androgen-dependent iridophore development (e.g. guppies^12^; *Rhodeus ocellatus ocellatus*^11^), the molecular mechanism underlying androgen-dependent iridophore development remained unclear. We indicated that androgen-dependent iridophore development is mediated by the Alkal2a-Ltk pathway in spotnape cardinalfish. Alkal-Ltk signaling that induces iridophore differentiation and proliferation was recently identified in zebrafish, in which defective mutants of Ltk or its ligand Alkal nullify iridophore development^27,45^, whereas hyperactive mutants of Ltk or those overexpressing Alkals exhibit ectopic iridophores^26,34^. Our androgen treatment analysis both *in vivo* and *in vitro* indicated that *alkal2a* is upregulated by androgens, and inhibitor treatment suggested that androgen-induced iridophore development is mediated by the Alkal2a-Ltk pathway (Figure 4). This finding is the first to demonstrate that the Alkal-Ltk pathway is also activated and is essential for androgen-dependent iridophore development.

In conclusion, we found that mouthbrooding cardinalfishes exhibit a whiter lower jaw, which may serve to conceal egg coloration during mouthbrooding. This trait may represent a repurposing of androgen-dependent body coloration, most of which have been reported to serve as conspicuous signals. Investigation of animals exhibiting unique traits such as mouthbrooding will provide us further interesting biological insights in the future.

## Materials and methods

### Animals

Spotnape cardinalfish (*Ostorhinchus notatus*) were caught by fishing in Manazuru city, Kanagawa Prefecture, Japan, unless otherwise mentioned. After they were caught, they were kept in a 200-400 L tank under breeding conditions, with a water temperature of 22±2°C, a 12-hour light/12-hour dark cycle. The fish were fed 1-2 times per day with hikari crest carnival (Kyorin, Hyogo, Japan) or Tetra Krill-E (Spectrum Brands Japan, Kanagawa, Japan). We used sexually mature fish with a body weight of 7.10-18.32 g.

Smaller spotnape cardinal fish were caught by hand net in Higashiizu, Kamo district, Shizuoka Prefecture. The body weights of a male and a female were 0.92 g and 1.09 g, respectively, with corresponding total lengths of 4.6 cm and 4.7 cm, respectively. Since the minimum size of a sexually mature individual was reported to be approximately 7.0 cm^25^, we regard these fish as immature. We also confirmed their immature gonads using a stereoscopic microscope. Fish with white ovarian outer membranes were considered females, while those with transparent testis-like structures were considered males.

*Ostorhinchus doederleini* (body weight: 15.11-23.72 g) were caught by fishing in Numazu city, Shizuoka Prefecture, Japan, during the breeding season in August 2022. *Ostorhinchus semilineatus* (body weight: 7.24-10.86 g) was caught using a hand net in Higashiizu, Kamo District, Shizuoka Prefecture, during the breeding season in August 2022. *Sphaeramia nematoptera* (body weight: 2.47-2.48 g) was purchased from a commercial source. The sex and maturity of each fish were determined by observing the gonads using a stereoscopic microscope.

All experiments were carried out according to protocols approved by the animal care and use committee of the University of Tokyo (Permission number: PH23-1).

### Seasonal sampling

Seasonal sampling of spotnape cardinalfish was conducted in Manazuru city, Kanagawa Prefecture, Japan. Fish were caught by fishing on 15 March, 21 April, and 14 June in 2023, and the water temperatures of the area around the sampling points were 16-17°C, 17-18°C, and 22-23°C, respectively (according to the Japan Meteorological Agency home page https://www.data.jma.go.jp/kaiyou/data/db/kaikyo/daily/sst_HQ.html). Blood samples were obtained immediately after collection. The observation of the lower jaw and measurement of the gonad-somatic index (GSI) were conducted on fish temporarily kept in a tank with the same water temperature as the field within 2 days. GSI was calculated as the ratio of gonad weight to whole body weight.

### Gross observation of the lower jaw and developing egg masses

After sexually mature spotnape cardinalfish were deeply anesthetized with 0.02% tricaine methanesulfonate (MS-222), photographs of each part of the body were taken. For photographs of the lower jaw with the mouth opening, tweezers were used to open the mouth. The parts taken in the photographs are shown in Figure S1A. The developing egg masses were collected from the male mouth during mouthbrooding in the breeding tank. The photographs were taken using digital single-lens reflex digital camera (ILCE-6300, SONY, Tokyo, Japan). Contrast and brightness were modified using Imaging Edge Desktop (SONY).

### Quantification of transparency of the lower jaw

After the fish were deeply anesthetized with 0.02% MS-222, the lower jaws were isolated and subjected to transparency quantification. For a control image without a sample, we captured photographs of a Petri dish filled with phosphate-buffered saline (PBS), the bottom of which was coated with silicone (KE-106, CAT-RG, Shin-etsu Chemical, Tokyo, Japan) to position the needles. For images of the lower jaws, we took photographs of the dish with the lower jaw immobilized by needles. A stereoscopic microscope (Leica M165 FC, Leica, Wetzlar, Germany) equipped with a 16-bit monochrome sCMOS camera, Zyla4.2PLUS (Oxford Instruments, Belfast, UK), was used for image acquisition. The photographs were captured under transmitted light with consistent light intensity using an A4 LED tracing table (ICHIREN, Hiroshima, Japan). For the analyses of the images, ImageJ (National Institutes of Health, Bethesda, MD) was used. The transparency was calculated by dividing the light intensity of the photograph with the sample by that of the photograph without the sample and then multiplying by 100 to obtain a percentage. For the calibration of the slight difference in the light intensity for each image acquisition, the average ratio of the intensities of the two peripheral areas of the sample photos and those of the corresponding control photos was used. The formula used for this calculation is as follows.

Transparency

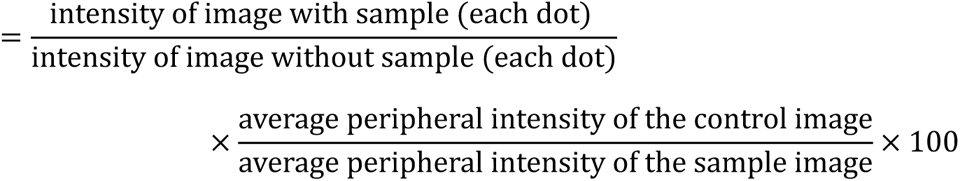

The area used for measuring transparency is described in Figures S1A-C.

### Artificial insertion of fertilized eggs into the mouth, which simulates assumed mouth breeding in each sex

The egg masses were collected by the same methods as described above from parental males at either 2 or 7 days into mouthbrooding. These egg masses were introduced to the mouths of other mature males or females after deep anesthetization by MS-222. Subsequently, the mouth was simply sutured to keep it closed and was photographed with a digital camera (ILCE-6300, SONY). Contrast and brightness were modified using Imaging Edge Desktop (SONY).

### Underwater photographs of the mouthbrooding males

Underwater photographs of the mouthbrooding males were captured in the breeding tank using an underwater camera (Tough TG-6, Olympus, Tokyo, Japan). The camera was remotely controlled via a smartphone application (OLYMPUS Image Share, Olympos) to minimize artificial effects on the brooding fish. Contrast and brightness were modified using OM Workspace (Olympus).

### Observation of the lower jaw and measurement of the iridophore-spreading area

After mature fish and fish subjected to MT-treatment experiments for 16 days were deeply anesthetized with 0.02% MS-222, the lower jaws were isolated. The lower jaws were immobilized on a Petri dish filled with PBS by pinning them to a black rubber sheet attached to the bottom of the dish. Note that the representative photograph in Figure 2 used transparent silicone instead of a black rubber sheet. Images of each part of the lower jaw were acquired with a WRAYCAM-NOA630B (WRAYMER, Osaka, Japan) mounted on a stereoscopic microscope (Olympus SZX16, Olympus). The areas subjected to observation are shown in Figures S1A, S1D, S1E.

For quantification of the area of iridophores, photographs were taken with consistent light intensity under reflected light. Since iridophores possess the property of reflecting light, the areas exhibiting light intensities greater than the threshold were regarded as the area of iridophores. The percentage of iridophores was calculated by dividing the area of iridophores by the total area subjected to quantification. The areas subjected to quantification are shown in Figures S1A-C.

### Histological observation of the lower jaw

After mature fish were deeply anesthetized with 0.02% MS-222, the lower jaws were isolated and fixed with 4% paraformaldehyde (PFA) in PBS. The fixed tissues were decalcified with 0.5 M EDTA (pH 8.0) and dehydrated in a graded ethanol series. Subsequently, the tissues were cleared with methyl benzoate and embedded in paraffin (Paraplast plus, Leica Biosystems, Wetzlar, Germany). The paraffin sections were cut at a thickness of 10 μm using a Leica Biocut 2035 (Leica Biosystems) and mounted on MAS-GP slide glass (Matsunami Glass, Osaka, Japan). For hematoxylin-eosin (HE) staining, the sections were deparaffinized with xylene and rehydrated in an ethanol series. After the sections were washed in distilled water, they were stained with Mayer’s hematoxylin for 10 minutes, followed by rinsing in tap water for 10 minutes. Then, the sections were stained with eosin solution for 1 min and washed in distilled water. The stained sections were dehydrated in a graded ethanol series, cleared with xylene, and then mounted with Permount (Thermo Fisher Scientific, Waltham, MA). Histological observations and polarized observations were conducted using an OLYMPUS BX53 (Olympus) instrument equipped with polarized optics and a WRAYCAM-NOA630B camera (WRAYMER).

### Measurement of plasma 11-KT levels

Blood samples were obtained from seasonally collected fish using a 1 mL syringe with 25 G needles coated with heparin sodium. Then, the samples were centrifuged at 3,000×g, and the supernatant was collected as a plasma sample. Five-microliter plasma samples were subjected to steroid extraction. To each plasma sample, 20 μL of diethyl ether was added, mixed with a vortex mixer, and allowed to stand to separate the supernatant layer of diethyl ether. After collecting the supernatant layer, the same procedure was repeated three more times to increase the yield. The collected supernatant was dried under a stream of nitrogen at 37°C. The extract was dissolved in 100 μL of EIA buffer (Cayman Chemical, Ann Arbor, MI). The 11-keto testosterone concentration in the extract was measured using an 11-keto testosterone ELISA Kit (Cayman Chemical) according to the manufacturer’s instructions. The absorbance was measured with a microplate reader (Multiskan FC, Thermo Fisher Scientific). The concentrations of 11-KT were calculated using Arigo Biolaboratories GainData (https://www.arigobio.com/elisa-analysis).

### Administration of androgen

A stock solution of 1 mg/mL 17α-methyltestosterone (MT) was prepared by dissolving MT (Tokyo Chemical Industry, Tokyo, Japan) in EtOH. The solution was directly added to the breeding water at a final concentration of 33 nM. The concentration of MT was determined based on previous reports^46–48^. The same amount of EtOH as the solvent was added to the control group. The tanks used for treatment were filled with 4 L of seawater, and one fish was treated per tank. The breeding water was maintained at a water temperature of 22±1°C, and the water was fully changed every other day. Fresh MT solution was added every day.

### RNA-seq analysis of MT-treated fish

The musculus intermandibularis of MT-treated fish was isolated and subjected to RNA-seq analysis. RNA extraction was conducted using a Fast Gene RNA Basic Kit (Nippon Genetics, Tokyo, Japan) according to the manufacturer’s protocol. The quality of the extracted RNA was confirmed by a 4200 TapeStation (Agilent Technology, Santa Clara, CA). The extracted RNA was purified by the NEBNext Poly(A) mRNA Magnetic Isolation Module (New England BioLabs, Ipswich, MA). The libraries for the Illumina sequencer (paired-end 150 bp) were prepared using the NEBNext Ultra II RNA Library Prep Kit for Illumina (New England BioLabs) according to the manufacturer’s protocol. The sequencing of the libraries was outsourced to a commercial sequence service (Nippon Genetics) using an Illumina NovaSeq 6000 (Illumina, San Diego, CA).

The low-quality reads and adaptor sequences were trimmed from the raw sequence data by fastp, version 0.22.0^49^, with the -q 30 option. The trimmed reads were subjected to *de novo* assembly using Trinity, version 2.15.0^50^, with default parameters. For *de novo* assembly of the transcriptome data of the lower jaw, we also used sequence data from various experiments involving a mature male, a mature female, and 11-KT- or MT-treated females. The trimmed reads were mapped to assembled contigs, and the expression of each contig was quantified by Salmon, version 0.10.0^51^, with default parameters. The transcript-level data were integrated to the gene level by using tximport version 1.22.0^52^. Identification of differentially expressed genes (DEGs) was conducted using edgeR, version 3.36.0^53^, with the filterByExpr option. Genes with a false discovery rate (FDR) less than 0.05 and log2-fold fold change (log2FC) less than -1 or greater than 1 were identified as DEGs. For annotation of identified DEGs, the sequences of DEGs were extracted from assembled transcriptome data by seqkit, version 2.2.0^54^. The isoforms from the same gene in the extracted sequences were clustered using cd-hit-est, version 4.8.1^55^. Then, the sequences of the DEGs were subjected to a homology search using BLAST, version 2.6.0+^56^, to the annotation data of a closely related species, *Sphaeramia orbicularis* (RefSeqID: GCF_902148855.1). The genes annotated as iridophore-related genes were subjected to manual annotation by phylogenetic analysis using ORTHOSCOPE^57^.

### Reverse transcription quantitative PCR (RT‒qPCR)

After mature males and females were deeply anesthetized with 0.02% MS-222, connective tissues and musculus intermandibularis in the lower jaws, upper jaws, and abdominal skins were isolated and subjected to RNA extraction using ISOGEN (Nippon Gene, Toyama, Japan) according to the manufacturer’s protocol. Then, the extracted RNA was treated with DNase Turbo (Thermo Fisher Scientific) to remove genomic DNA. Complementary DNA (cDNA) was synthesized using the PrimeScript RT Reagent Kit (Takara Bio, Shiga, Japan).

For the MT-treated and corresponding control fish, connective tissues and musculus intermandibularis in the lower jaws and abdominal skins were isolated and subjected to RNA extraction using a Fast Gene RNA Basic Kit (Nippon Genetics) with DNaseI treatment. The extracted RNA was reverse-transcribed using the PrimeScript RT Reagent Kit (Takara Bio).

RT‒qPCR was conducted by using a KAPA SYBR Fast qPCR kit (Nippon Genetics) with a LightCycler 480 II system (Roche Applied Science, Penzberg, Germany). The PCR reactionwas conducted under the following conditions: 95°C for 5 minutes; 45 cycles of 95°C for 10 seconds, 60°C for 10 seconds, and 72°C for 10 seconds. The PCR products were subjected to melting curve analysis for verification. For normalization, the housekeeping gene *β-actin* (*actb*) was used. Note that the primers for each reaction were designed based on the predicted mRNA sequence assembled by Trinity. To obtain sequences of target genes, candidate transcript contigs were searched from our transcriptome data using BLAST, version 2.6.0+^56^, with orthologs of other species as queries. Then, to confirm their orthologies, the candidate contigs were subjected to phylogenetic analysis using ORTHOSCOPE^47^. The sequences of the primers used in the experiments are shown in Table S1.

### Reverse transcription PCR (RT‒PCR)

A mature male was deeply anesthetized with 0.02% MS-222. Connective tissue and musculus intermandibularis in the lower jaw, abdominal skin, and testis were isolated and subjected to RNA extraction using ng ISOGEN (Nippon Gene). Then, the RNA was treated with Turbo DNase (Thermo Fisher Scientific). cDNA was synthesized as described above using the PrimeScript RT Reagent Kit (Takara Bio).

RT‒PCR was conducted with a KAPA HiFi Hotstart ReadyMix PCR Kit (Nippon Genetics). The PCR reaction was conducted under the following conditions: 98°C for 4 minutes; 30 cycles of 98°C for 20 seconds, 62°C for 15 seconds, and 72°C for 15 seconds; and a final extension at 72°C for 1 minute. The PCR products were visualized by gel electrophoresis in a 1.5% agarose gel. The primers used in this experiment are shown in Table S1.

### cDNA cloning and *in situ* hybridization

cDNAs of *pnp4a*, *alkal2a*, and *ara* were cloned to make mRNA probes for *in situ* hybridization. Their cDNA sequences were amplified by RT‒PCR as described above. Then, the PCR products were ligated into the pGEM-T Easy Vector (Promega, Madison, WI), and the sequences were confirmed by Sanger sequencing. Digoxigenin (DIG)-labeled probes were synthesized with a DIG RNA Labeling Kit (Roche Applied Science). The paraffin sections were made as described above and subjected to *in situ* hybridization. *In situ* hybridization was conducted as previously reported^58^. We confirmed that signals were detected in sections hybridized with the antisense probe but were not detected in sections hybridized with the sense probe (Figure S9). The sections were observed with an OLYMPUS BX53 (Olympus) instrument equipped with a WRAYCAM-NOA630B camera (WRAYMER). The primers used in this experiment are shown in Table S1.

### Alignment of Alkal2a and prediction of domain structure

The amino acid sequence predicted from the cDNA sequence obtained by cloning, as described above, was subjected to multiple protein alignment using CrustalW in MEGA-11^59^. The alignment was visualized by Jalview^60^. The following amino acid sequences were used in this analysis: Alkal2a-NP_001410815.1 (*Danio rerio*), XP_011489918.1 (*Oryzias latipes*), Alkal2-NP_001002919.2 (*Homo sapiens*), NP_001153215.1 (*Mus musculus*), and XP_004914540.1 (*Xenopus tropicalis*). Then, the alignment was subjected to domain structure prediction using NCBI Batch CD-search^61^.

### *In vitro* culture of lower jaws

After mature males and females were deeply anesthetized with 0.02% MS-222, the lower jaws were isolated. Connective tissues and the anterior and posterior regions of the musculus intermandibularis were isolated from the lower jaws using a sterilized razor under a stereoscopic microscope. Connective tissue was obtained from the right and left sides of the body, resulting in two identical tissues from each fish. The musculus intermandibularis was cut at midline, resulting in two identical tissues from each fish. The surfaces of these tissues were sterilized by soaking in 70% EtOH for a few seconds, followed by washing in sterilized PBS. Then, the tissues were cultured at 22°C in a 12-well plate filled with 2 mL of medium. The medium was fully changed every 3 days.

The basic medium was composed of Leibovitz L-15 medium (FUJIFILM Wako Pure Chemical Corporation, Osaka, Japan) supplemented with 10% heat-inactivated fetal bovine serum (FBS), 1% 100× penicillin‒streptomycin, 1% 100× amphotericin B, and 0.1% dimethyl sulfoxide (DMSO) as a solvent for drugs. For experiment 1, tissue from the fish was cultured in basic medium, while the other tissue was cultured in basic medium supplemented with 33 nM MT. For experiment 2, tissue from the fish was cultured in basic medium supplemented with 33 nM MT, while the other tissue was cultured in basic medium supplemented with 33 nM MT in addition to 100 nM of inihibitor of Leulocyte tyrosine kinase (Ltk): TAE-684 (TargetMol, Boston, MA). After 8 days of treatment, RNA was extracted using a Fast Gene RNA Basic Kit (Nippon Genetics) with DNaseI treatment. Then, qRT‒PCR was conducted as described above.

### Observation of sexual behavior

Spotnape cardinalfish form male/female pairs during the breeding season and are kept together for a few days to a few weeks, after which they spawn^62,63^. We recorded their sexual behavior using a video camera (HDR-CX420, SONY) in breeding tanks for more than five minutes after the pair had formed. In each video, we recorded their sexual behavior for five minutes after they formed pairs using the Excel macro Ethogramer^64^. The definitions of each behavior are as follows. Display with warping: The behavior of approaching the partner with body warping. Circling: The behavior of circling above or below its partner. Attack to other fish: The behavior of chasing other fish that come close to the pair. Note that the tank used for observing pair 1 was filled with approximately 150 L of seawater containing 15 fish, with a sex ratio of approximately 1:1. The behaviors of pairs 2 and 3 were observed in the same tank, which was filled with approximately 150 L of seawater containing 15 fish, with a sex ratio of approximately 1:1. The tank used for observing pair 4 was filled with approximately 400 L of seawater containing 30 fish, with a sex ratio of approximately 5:1 male to female.

### Statistical analysis

For comparisons between non-paired two groups, the data were analyzed with the Wilcoxon rank sum test using R, version 4.1.2. For comparisons between paired two groups in *in vitro* culture experiments, the data were analyzed with the Wilcoxon signed-rank test using R, version 4.1.2. For comparison among more than three groups, the data were processed with the Steel–Dwass test using Kyplot 6.0 software (Kyence, Tokyo, Japan). Graphs were drawn with R, version 4.1.2. A P value less than 0.05 was considered to indicate statistical significance.

## Acknowledgments

We thank Mr. Kosei Inaba (Tokyo University of Marine Science and Technology) for providing *Ostorhinchus semilineatus* and for his cooperation with sample collection. We are grateful to Mr. Masahiro Takano (Tokyo University of Marine Science and Technology), Drs. Soma Tomihara (Nagahama Institute of Bio-Science and Technology), Drs. Ayaka Fukuda, Mr. Katsumi Yokota, Ms. Mana Yamakawa, Mr. Shun Kenny Uehara, Mr. Kohei Ochiai, and Mr. Manabu Yamzaki (The University of Tokyo) for their cooperation with sample collection. We also thank Drs. Satoshi Kawato (National Institute of Infectious Diseases) for his helpful comments on RNA-Seq analysis. We are grateful to Mr. Koya Shimoyama (The University of Tokyo) for helpful advice on ELISA and RNA-Seq analysis as well as to Drs. Naotaka Aburatani for his guidance on paraffin sectioning. We are grateful to Vesper Studio (Tokyo, Japan) for the preparation of the schematic illustration.

## Author contributions

Conceptualization: H.I. and S.K

Investigation: H.I. and S.K.

Supervision: S.K.

Writing—original draft: H.I. and S.K.

Writing—review & editing: H.I. and S.K.

Project administration: S.K.

Funding acquisition: S.K.

## Data and materials availability

- All raw sequencing data generated in this study have been submitted to the NCBI BioProject database (https://www.ncbi.nlm.nih.gov/bioproject/) under accession number PRJDB18055.
- This paper does not report original code.
- All other data reported in this paper is available from the lead contact upon request from the lead contact.

## Declaration of generative AI and AI-assisted technologies

The draft written by the authors was proofread by AI-powered scientific writing assistant, Curie (https://www.aje.com/curie/).

## Supplementary Materials

Figures S1 to S10

Table S1

