## Supplemental figures and table for "Inconspicuous breeding coloration to conceal eggs during mouthbrooding in male cardinalfish"

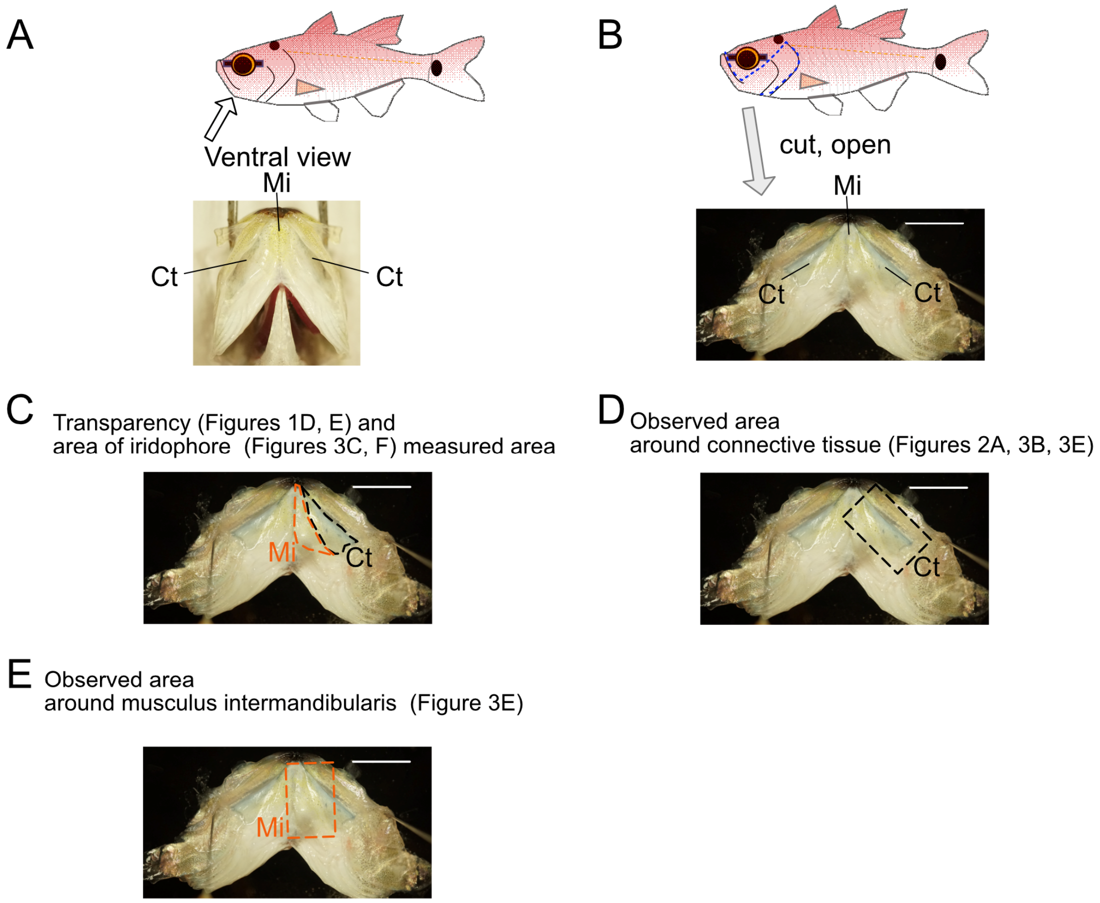


**Figure S1 Description of the lower jaw region used in each observation and analysis.**

(A) Ventral view of the lower jaw with the mouth open. The photograph was taken in the area marked with white arrows. (B) The isolated lower jaw cut out by the blue dashed line on both sides of the body. After the incision was made, the lower jaw was opened so that the outside of the cavity was up. (C) Transparency (Figures 1D and 1E) and area of the iridophore (Figures 3C and 3F) measured region. The black dashed line indicates the region measured as Ct. The orange dashed line indicates the region measured as Mi. (D) The black dashed line indicates the observed region around Ct (Figures 2A, 3B, and 3E). (E) The orange dashed line indicates the observed region around Mi (Figure 3E). Ct, connective tissue; Mi, musculus intermandibularis. Scale bar, 10 mm.

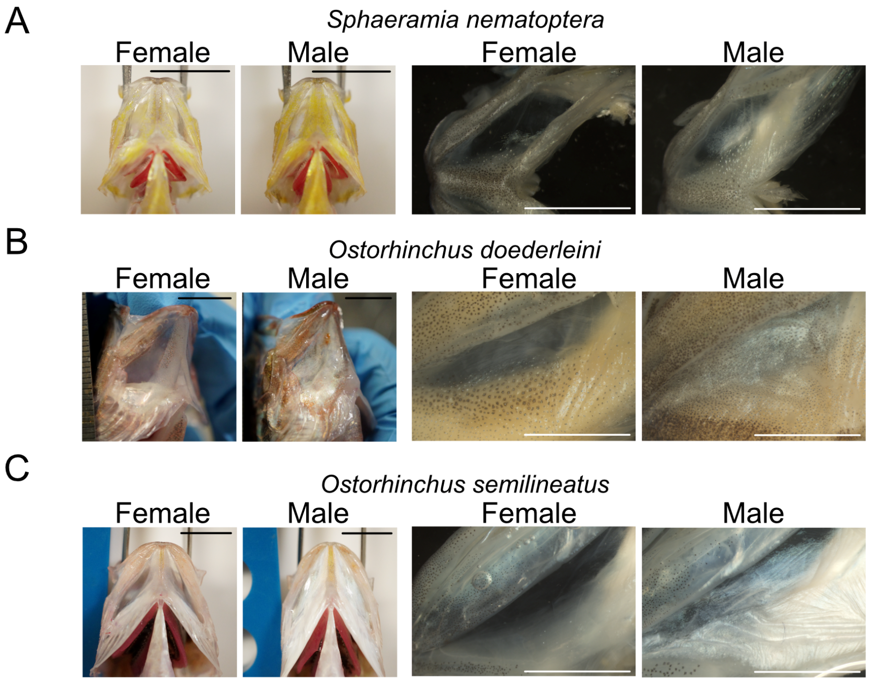


**Figure S2 The lower jaws of other male Apogoninae fishes also contain iridophores.**

(A) *Sphaeramia nematoptera*, (B) *Ostorhinchus doederleini*, and (C) *Ostorhinchus semilineatus*. Note that the right photograph shows an enlarged view of samples after fixation with 4% PFA. Scale bar, 10 mm.

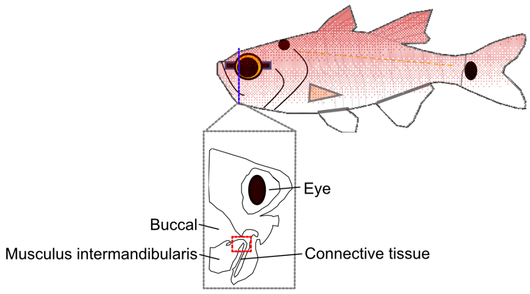


**Figure S3. Schematic illustration of the sectioning site.**

The frontal illustration surrounded by a gray dashed line indicates the frontal plane of the blue dashed line on the fish illustration. The area surrounded by the red dashed line indicates the area observed in Figures 2B, S3B, and 4I.

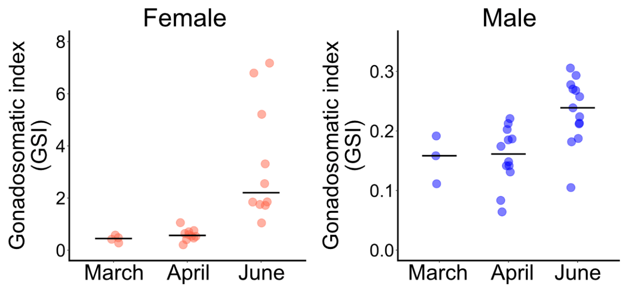


**Figure S4 Seasonal changes in the gonadosomatic index (GSI) of males and females.**

Gonadal size increases as the breeding season approaches. Bars indicate the medians. females in March: n = 4 males in March: n = 3; females in April: n = 10; males in April: n = 12; females in June: n = 10; males in June: n = 13. Due to the small number of samples in March, statistical test has not been performed.

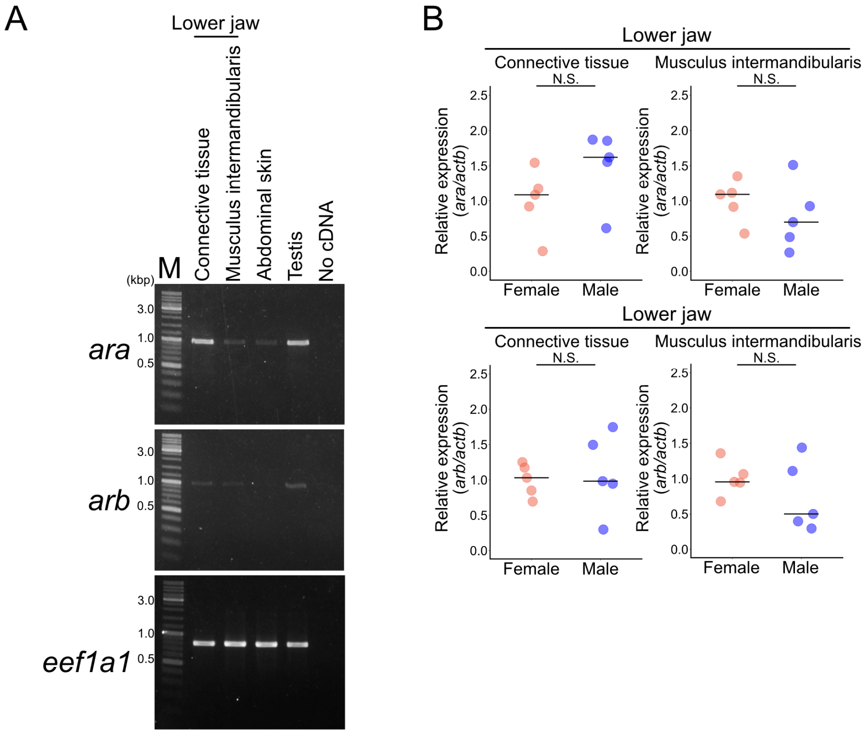


**Figure S5 The genes encoding androgen receptors are expressed in both male and female lower jaws.**

(A) RT–PCR for *androgen receptor a* (*ara*) and *b* (*arb*) in male fish. The expression of *ara* was detected in the connective tissue and musculus intermandibularis in the lower jaw. A small amount of the expression of *arb* was detected in the connective tissue and musculus intermandibularis in the lower jaw. (B) The expression levels of *ara* and *arb* in connective tissue and musculus intermandibularis in the lower jaw were not significantly different between females and males. Bars indicate the medians. N.S., not significant, Wilcoxon rank sum test; n = 5.

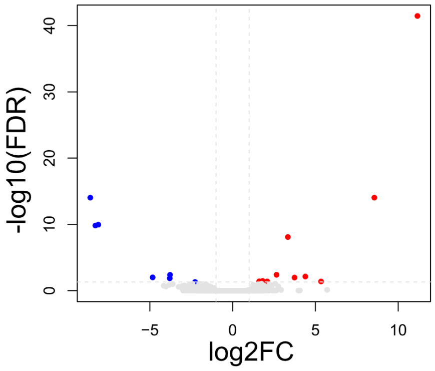


**Figure S6 Volcano plot of differentially expressed genes (DEGs) from RNA-seq analysis of musculus intermandibularis in the lower jaw between EtOH-treated females and 33 nM methyl testosterone-treated females on day 3.**

Positive log2 fold-change (FC) values correspond to increased expression in females treated with 33 nM methyl testosterone. The vertical lines indicate a log2FC threshold of 1, while the horizontal line indicates a -log10 false discovery rate (FDR) threshold of -log10(0.05).

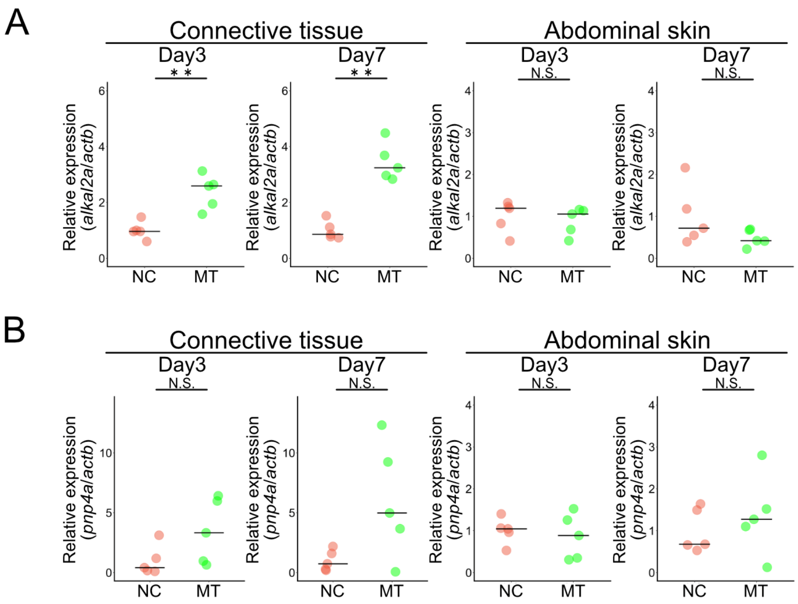


**Figure S7 Expression of *alkal2a* is upregulated by androgen prior to iridophore development, specifically in the lower jaw.**

(A) The expression level of *alkal2a* of 33 nM methyl testosterone-treated females (MT) on days 3 and 7 is significantly greater than that of EtOH-treated females (NC) in the connective tissue in the lower jaw, whereas it is not significantly different in other tissues, such as the abdominal skin. Bars indicate the medians. **, $p < 0.01$, Wilcoxon rank sum test; $n = 5$. (B) The expression levels of *pnp4a* in the connective tissue in the lower jaw and abdominal skin does not differ between 33 nM methyl testosterone-treated females (MT) and EtOH-treated females (NC) on days 3 and 7. Bars indicate the medians. N.S., not significant, Wilcoxon rank sum test; $n = 5$.

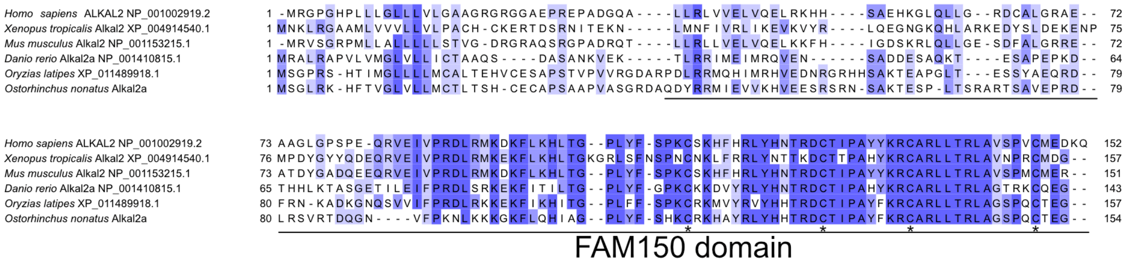


**Figure S8 Alignment of Alkal2 proteins.**

The underline indicates the FAM150 domain predicted in Alkal2a of spotnape cardinalfish. Asterisks indicate four conserved cysteines essential for maintaining structural integrity.

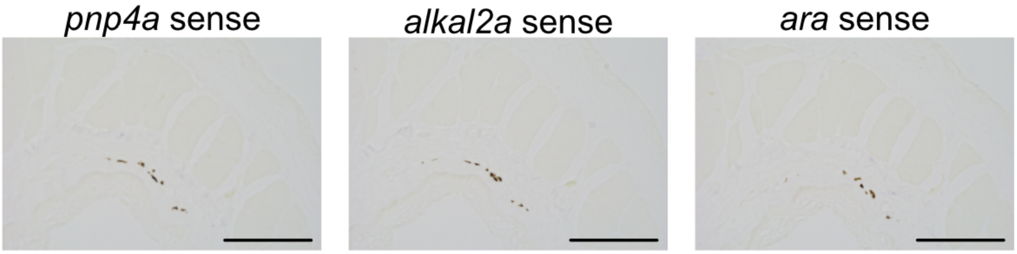


**Figure S9 Confirmation of the specificity of mRNA probes using sense probes.**

No purple signals are detected with the control sense probe. Scale bar, 100 µm.

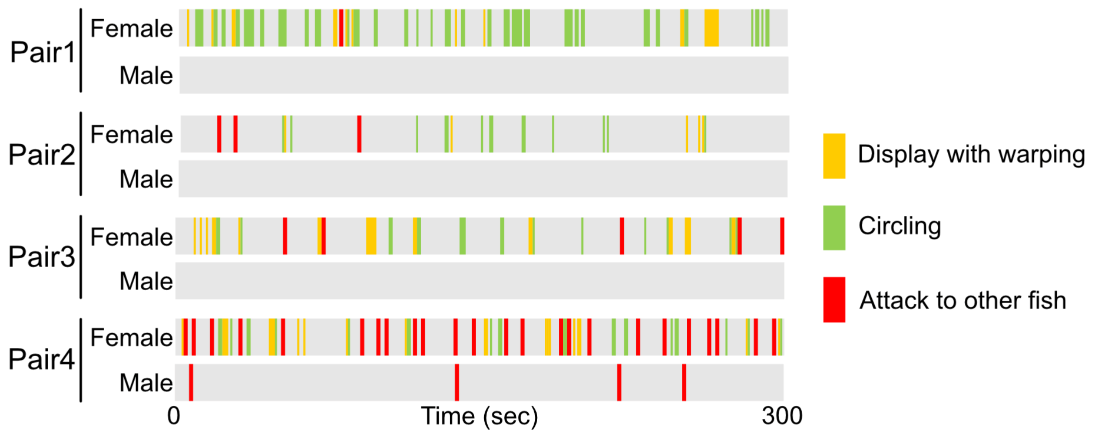


**Figure S10 Time-course data of 5-min analyses of sexual behavior are shown as raster plots.** The yellow, green, and red bands indicate the timing and duration of display with warping, circling and attack to other fish, respectively.

Table S1 List of primer sequences.

| Primer used for RT-qPCR | |
|---|---|
| Name | Sequence (5' to 3') |
| actb_F | AGCACAGTGTTGGCGTACAG |
| actb_R | CCTTCCTTCCTCGGTATGGA |
| pnp4a_F | CTGGTATTCGGAGAGTTGAAGG |
| pnp4a_R | AACTGGGAACGTTGTCTTACAGAG |
| ara_F | GCTCTATGCTCTACTTTGCTCCA |
| ara_R | ACCTTCAGCATACAAAACCTCTG |
| arb_F | GAAGAACTCCATGGGGAACA |
| arb_R | GAAGGCAGAGGTGGGAGAGT |
| alkal2a_F | AAAGATGTGCACGGCTTCTC |
| alkal2a_R | GACAACTCGTATTTGGCTTCAAC |

| Primer used for RTPCR | |
|---|---|
| Name | Sequence (5' to 3') |
| ara_F | GAGTCTGATCCGTTGGATAC |
| ara_R | ACAGGTGGTTCTGCTTACCT |
| arb_F | ACCGTGTCCTGCTCTTATGG |
| arb_R | TGAAACCTGGGAGTCCTTTG |
| eef1a1_F | GCCTACATTAAGAAGATCGGTTACA |
| eef1a1_R | TCTTCTCCACTGACTTGATAACACC |

| Primer used for cloning for in situ hybridization | |
|---|---|
| Name | Sequence (5' to 3') |
| pnp4a_F | GCTGATTGGCTGATGTCTCA |
| pnp4a_R | GGCAAAACCAATCAACATGA |
| alkal2a_F | GCTTCCTGTAACCGCGTATC |
| alkal2a_R | TGTGTGCTACCCTTCTGTGC |
| ara_F | GAGTCTGATCCGTTGGATAC |
| ara_R | ACAGGTGGTTCTGCTTACCT |
